# CORE CONSERVED TRANSCRIPTIONAL REGULATORY NETWORKS DEFINE THE INVASIVE TROPHOBLAST CELL LINEAGE

**DOI:** 10.1101/2023.03.30.534962

**Authors:** Ha T.H. Vu, Regan L. Scott, Khursheed Iqbal, Michael J. Soares, Geetu Tuteja

## Abstract

The invasive trophoblast cell lineage in rat and human share crucial responsibilities in establishing the uterine-placental interface of the hemochorial placenta. These observations have led to the rat becoming an especially useful animal model to study hemochorial placentation. However, our understanding of similarities or differences between regulatory mechanisms governing rat and human invasive trophoblast cell populations is limited. In this study, we generated single-nucleus (sn) ATAC-seq data from gestation day (gd) 15.5 and 19.5 rat uterine-placental interface tissues and integrated the data with single-cell RNA-seq data generated at the same stages. We determined the chromatin accessibility profiles of invasive trophoblast, natural killer, macrophage, endothelial, and smooth muscle cells, and compared invasive trophoblast chromatin accessibility to extravillous trophoblast (EVT) cell accessibility. In comparing chromatin accessibility profiles between species, we found similarities in patterns of gene regulation and groups of motifs enriched in accessible regions. Finally, we identified a conserved gene regulatory network in invasive trophoblast cells. Our data, findings and analysis will facilitate future studies investigating regulatory mechanisms essential for the invasive trophoblast cell lineage.

## INTRODUCTION

Hemochorial placentation is a reproductive strategy utilized by some mammals, including the mouse, rat, and human [1]. This type of placentation involves establishment of a uterine-placental interface characterized by trophoblast cells of extraembryonic origin breaching the maternal vasculature [1]. Trophoblast cells are the parenchymal cells of the placenta [2–4]. Their origins can be traced to the trophectoderm of the early embryo and the initial cell differentiation event during embryogenesis [5,6]. Trophoblast cells differentiate into a range of specialized lineages [3–5]. Among the specialized trophoblast cell lineages are invasive trophoblast (generic term) or extravillous trophoblast (**EVT**, human/primate specific term). These cells exit the placenta and enter the uterine compartment where they transform the vasculature and immune environment into a structure ensuring placental and fetal viability and growth [3,4,7]. Failures in invasive trophoblast/EVT cell differentiation and function result in a range of pregnancy diseases such as preeclampsia, intrauterine growth restriction, and preterm birth [8,9]. Deep trophoblast cell invasion and uterine transformation are characteristic features of rat and human placentation sites [10–13]. Identification of potential regulatory mechanisms controlling cellular constituents of the rodent and human uterine-placental interface have emerged from single-cell RNA-sequencing (**scRNA-seq**) [14–20]. Conserved sets of transcripts have been identified in rat invasive trophoblast and human EVT cells [20]. These insights have led to the identification of candidate regulators of invasive trophoblast and EVT cell lineages and dissection of their biological relevance using trophoblast stem (**TS**) cells and rat models [21–23]. Such experimentation has advanced the field but on its own is an inefficient strategy for defining gene regulatory networks driving invasive trophoblast/EVT cell lineage development and function.

Gene regulatory networks can be accessed through genome-wide analysis of the chromatin landscape [24–27]. Indeed, insights into the hierarchical regulation of rodent and human trophoblast cell development have been achieved through deep sequencing of histone modifications defining gene activation and repression states [28–36]. The integration of transcriptome and chromatin accessibility datasets has also been used as an effective tool to elucidate gene regulatory networks in trophoblast tissue and cells [37,38].

In this report, we interrogated the chromatin landscape of invasive trophoblast cells isolated from the uterine-placental interface of the rat using single-nucleus assay for transposase-accessible chromatin-sequencing (**snATAC-seq**). These datasets were integrated with scRNA-seq datasets from rat and human invasive trophoblast/EVT cells [20], as well as ATAC-seq from EVT cells [39], to identify conserved gene regulatory networks controlling the invasive trophoblast cell lineage.

## RESULTS

### Identification of chromatin accessibility profiles in cell types of the rat uterine-placental interface

We generated snATAC-seq profiles from gestation day (**gd**) 15.5 and 19.5 uterine-placental interface tissue of the rat to determine chromatin accessibility of its cellular constituents. These datasets were integrated with scRNA-seq profiles obtained from the same tissues [20].

Following quality control and preprocessing (**Figs. S1 and S2**), we obtained 25,321 and 14,388 high quality nuclei in the gd 15.5 and gd 19.5 samples, respectively (**Table S1**). Next, snATAC-seq data was integrated with scRNA-seq data [20] to identify cell populations based on the relationship between accessibility and gene expression profiles [40] (**Fig. 1A**). Clusters and chromatin accessibility profiles of invasive trophoblast, natural killer, macrophage, endothelial, and smooth muscle cells were identified (**Table S1**).

**Fig. 1.**
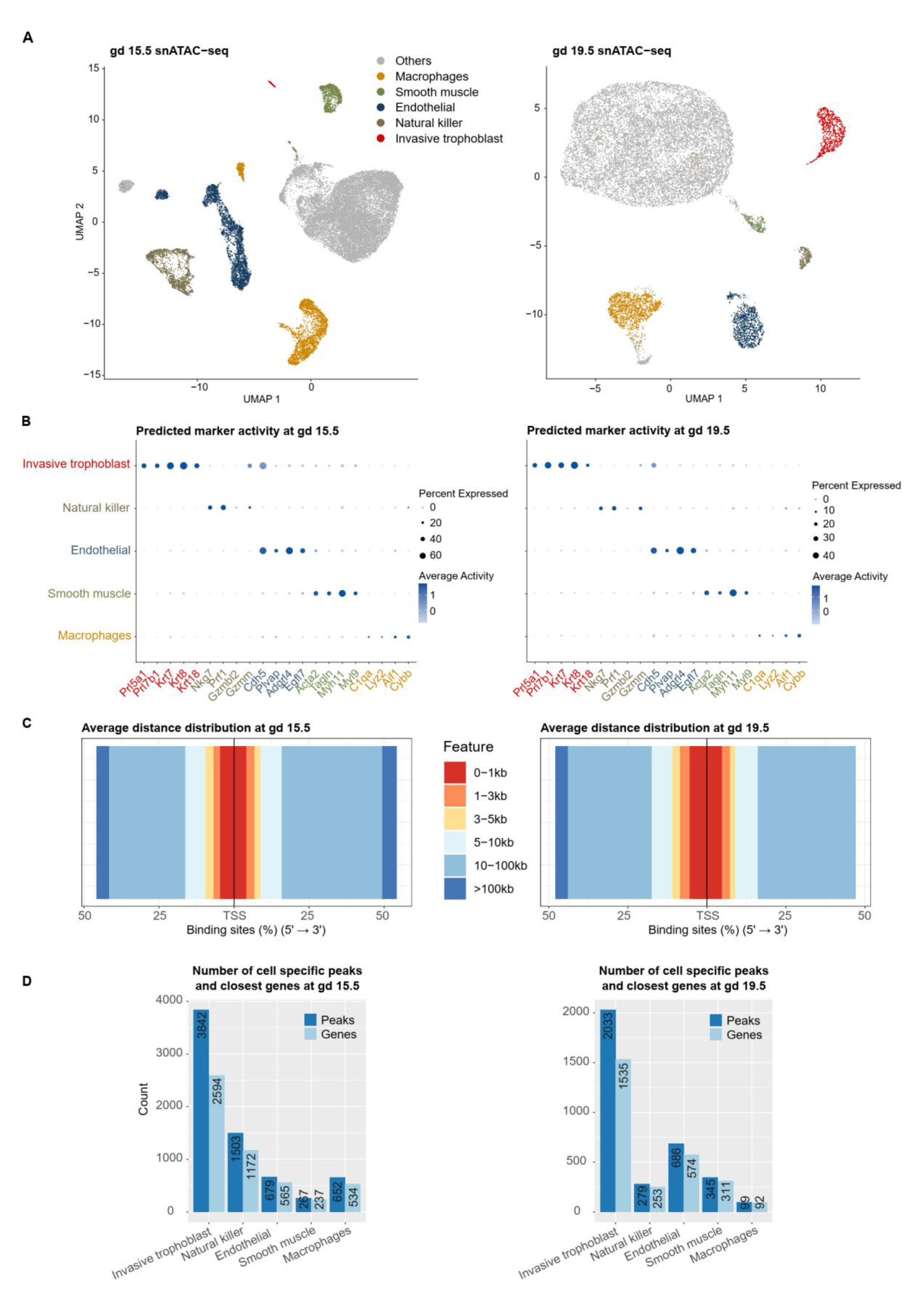
Chromatin accessibility profiles of cell populations at the uterine-placental interface. **A**) UMAP of snATAC-seq profiles at gestation day (**gd**) 15.5 and 19.5 showing cell identities obtained by transferring labels from scRNA-seq data. **B**) Dot plots showing known markers of cell types generally have higher accessibility within 2,000 base pairs (**bp**) of the transcription start sites (**TSS**) and in a higher percent of nuclei than in other cell populations. Dot sizes correspond to the percent of nuclei in each cell population that were open around the TSS; colors correspond to the levels of predicted gene activity. **C**) Stack bar plots showing that cell type-specific open chromatin peaks were most often distal to the TSS. For distribution of distances for each individual cell type see **Fig. S4. D**) Bar plots showing the number of open chromatin peaks specific to a cell population, and the number of nearest genes to cell-specific open chromatin peaks. Cell specific open chromatin peaks are open chromatin peaks differentially accessible in the cell population compared to all other cell populations (adjusted p-value ≤ 0.05, average log_2_(fold change) ≥ log_2_(1.5)).

These analyses are based on an assumption that there is a significant correlation between gene expression level (scRNA-seq data) and chromatin accessibility (snATAC-seq data) [40]. Therefore, as a quality control step for the snATAC-seq cluster labeling, we calculated the Spearman correlation between gene expression and chromatin accessibility profiles. We obtained moderate but significant correlations (0.44≤ρ≤0.54, p-value<2.2e-16) in all cell populations (**Fig. S3**), which agrees with previous studies done at the both single cell and tissue levels [38,41,42]. Moreover, we observed that established marker gene expression for each cell population are generally more accessible in the respective cell population (**Fig. 1B**), demonstrating we have obtained high quality clustering and cluster annotation.

We further performed differential accessibility analysis at both gestation days to identify the most accessible peaks in each cell type (defined as cell type-specific peaks). The distance distribution of cell type-specific peaks to the nearest gene transcription start site (**TSS**) showed, that in general, most of the cell type-specific peaks are distal to the TSS (>5 kb) (81.65% at gd 15.5, and 75.16% at gd 19.5) (**Fig. 1C, Fig. S4**). Moreover, we observed that the invasive trophoblast cell population had the highest number of cell type-specific peaks of the major cell types analyzed, despite being of less abundance than some other cell types (**Fig. 1D**). Related to this, the invasive trophoblast cell population had the most gene-associated accessible chromatin among the cell types identified.

### Identification of invasive trophoblast cell regulated genes using cell type-specific chromatin accessibility profiles

Following the observation that invasive trophoblast cells had the most cell type-specific peaks, we next checked the number of peaks associated with each gene at each gestation day. At both gestational timepoints, there were many genes associated with at least two open regions (808 and 349 genes at gd 15.5 and 19.5, respectively) (**Fig. 2A**). Next, we investigated the differences in expression levels of transcripts linked to 2, 3, 4, or 5 peaks using the average expression level obtained from the scRNA-seq data. In general, we observed an increasing trend of expression level when a transcript is associated with more peaks. Furthermore, we observed that while gd 15.5 expression levels were significantly different as the number of associated peaks increased, at gd 19.5, transcript expression profiles were not significantly different when more peaks were associated with a transcript after a cut-off of 3 (**Fig. S5**). Therefore, we partitioned transcripts into two groups for the next analyses: ≥3 peaks or <3 peaks. At both gestation days, genes with more than three peaks had significantly higher expression than genes with less than three peaks (p-value=7.029e-09 and 1.374e-08 at gd 15.5 and 19.5, respectively) (**Fig. 2B**), suggesting that, in general, genes with ≥3 trophoblast-specific peaks are more active within the cell population and could have important functional roles for trophoblast cells. However, there are notable exceptions, including *Prl7b1* (**Table S2**), which has <3 peaks, but whose expression is specific and among the highest in the invasive trophoblast cell lineage.

**Fig. 2.**
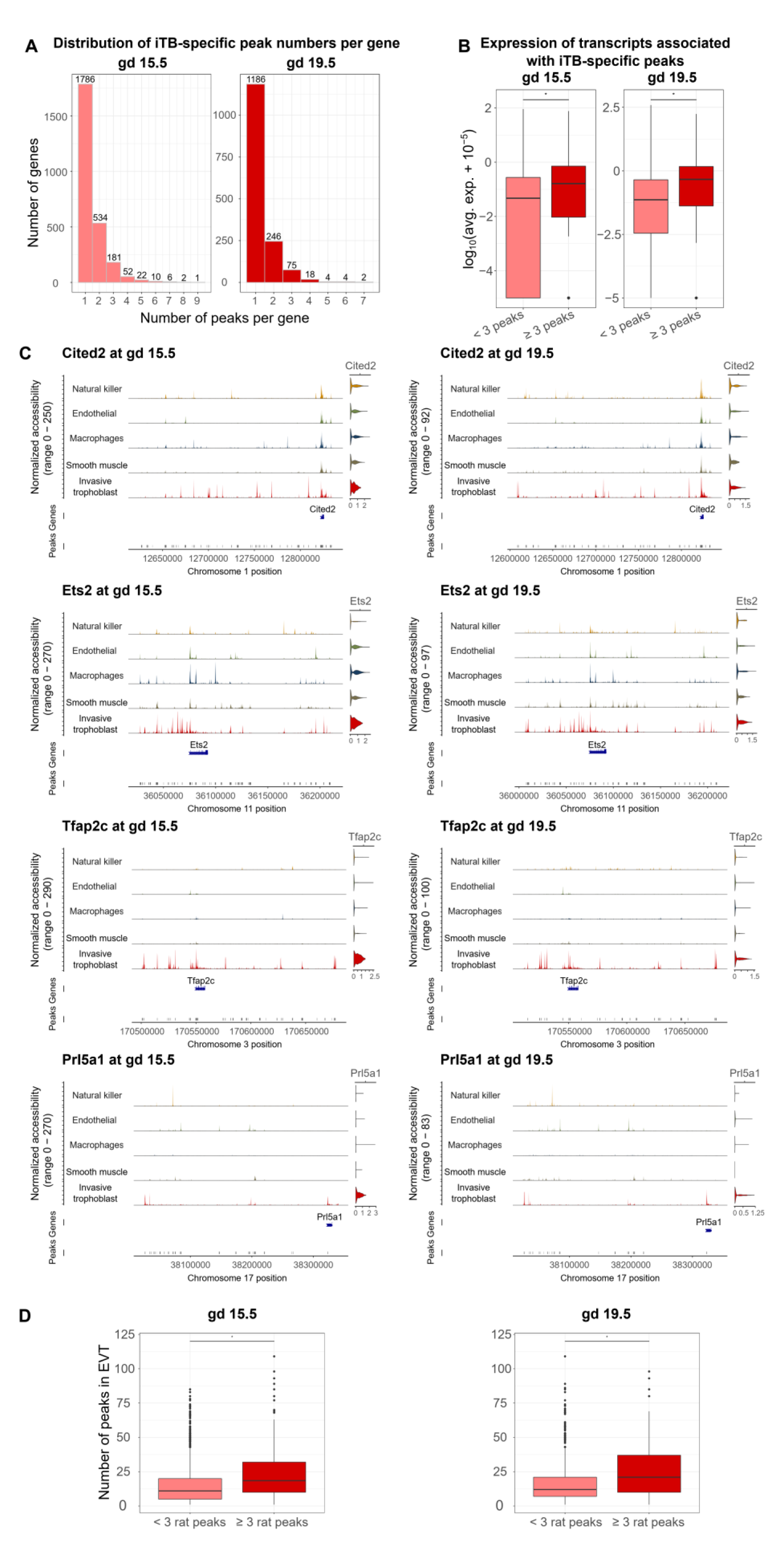
Analysis of chromatin accessibility profiles can identify regulatory regions for genes defining the invasive trophoblast cell population. **A**) Histograms of the number of invasive trophoblast-specific (**iTB-specific**) peaks per gene showing that many genes had ≥1 peaks. The x-axis shows the number of peaks per gene, and the y- axis shows the number of genes. **B**) Boxplots of transcript expression associated with iTB-specific peaks showing that genes with ≥3 peaks had significantly higher expression than genes with fewer than 3 peaks. Expression was plotted in a log_10_(average expression + 10^-5^) scale. **C**) Examples of iTB-specific genes with ≥3 associated peaks at both gd 15.5 and 19.5. For each subplot, the first section was composed of five tracks of normalized accessibility, corresponding to five cell types. The right-most column of the first section shows the predicted gene activity using chromatin accessibility within 2,000 bp of the TSS. The second and third section include two tracks corresponding to gene location and open chromatin peak locations, respectively. **D**) Boxplots of the number of conserved ATAC-seq peaks in EVT cells and rat invasive trophoblast cells. Rat genes with ≥3 invasive trophoblast cell-specific peaks had significantly more EVT cell ATAC- seq peaks than rat genes with <3 invasive trophoblast cell-specific peaks. ATAC-seq peaks in EVT cells were obtained from Varberg et al. [39]. Statistical analyses were performed using Wilcoxon rank sum tests at a significance level of 0.05.

In addition, we compared transcripts with ≥3 open regions to transcripts with invasive trophoblast cell cluster-specific expression (invasive trophoblast cell marker transcripts), previously determined from the scRNA-seq data [20] at each gestation day. At gd 15.5, 57 of the 274 genes with ≥3 peaks were also markers of the invasive trophoblast cell cluster (p-value=5.29e-08), and at gd 19.5, 39 of the 103 genes with ≥3 peaks were markers of the invasive trophoblast cell cluster (p-value=6.79e-06). These markers included genes with known trophoblast functions (*Tfap2c* [43–45], *Ets2* [46], and *Cited2* [23,47,48]), and genes known to be prominently expressed in invasive trophoblast cells (*Prl5a1* [49]) (**Fig. 2C**). Of note, while some of these markers (*Cited2* and *Ets2*) have similar activities around their promoter regions in all cell types, they had multiple associating peaks specific to the invasive trophoblast cell cluster.

To determine if transcripts that have multiple associated peaks in rat invasive trophoblast cells also possess multiple associated peaks in human EVT cells, we incorporated open regions (ATAC-seq peaks) identified in EVT cells into our analysis [39]. First, we associated the EVT cell open regions to genes. Then, we compared the number of EVT cell peaks associated with genes that have either ≥3 or <3 peaks in rat invasive trophoblast cells (**Table S2**). We observed that, at both time points, genes with ≥3 peaks in rat invasive trophoblast cells had significantly more peaks in human EVT cells than genes that had fewer than 3 peaks in rat invasive trophoblast cells (p-value<2.2e-16) (**Fig. 2D**).

### Identification of transcription factors (TFs) enriched in invasive trophoblast cell-specific peaks

To predict TFs that may be important for the invasive trophoblast cell population, we carried out motif enrichment analysis in the 1242 invasive trophoblast cell-specific peaks identified at both gestation days, hereafter referred to as common peaks (**Table S3**). Following the enrichment tests, filtering, and TF family grouping, we identified 11 TF families that were enriched in the common peaks, some of which have known roles in regulating trophoblast biology (**Fig. 3A, Table S3**). For example, TFAP2C motifs were enriched with the highest fold change in the common peaks. TFAP2C is a member of the AP-2 TF family and is a known regulator of the trophoblast cell lineage in both mouse and human [3,50,51]. We further confirmed the enrichment of the TFAP2C binding sites by comparing the rat open regions with TFAP2C motifs to TFAP2C chromatin immunoprecipitation (ChIP)-seq peaks from differentiated mouse TS cells [34]. We found that of the 439 rat peaks with TFAP2C motifs, 208 (47.38%) overlapped with TFAP2C peaks in differentiated mouse TS cells, which was significant (p-value=0.009). Additionally, all 11 TF families enriched in the rat peaks were enriched in EVT cell ATAC-seq peaks [39] (**Table S3**). These comparisons provide evidence for the validity of the computationally based binding site predictions.

**Fig. 3.**
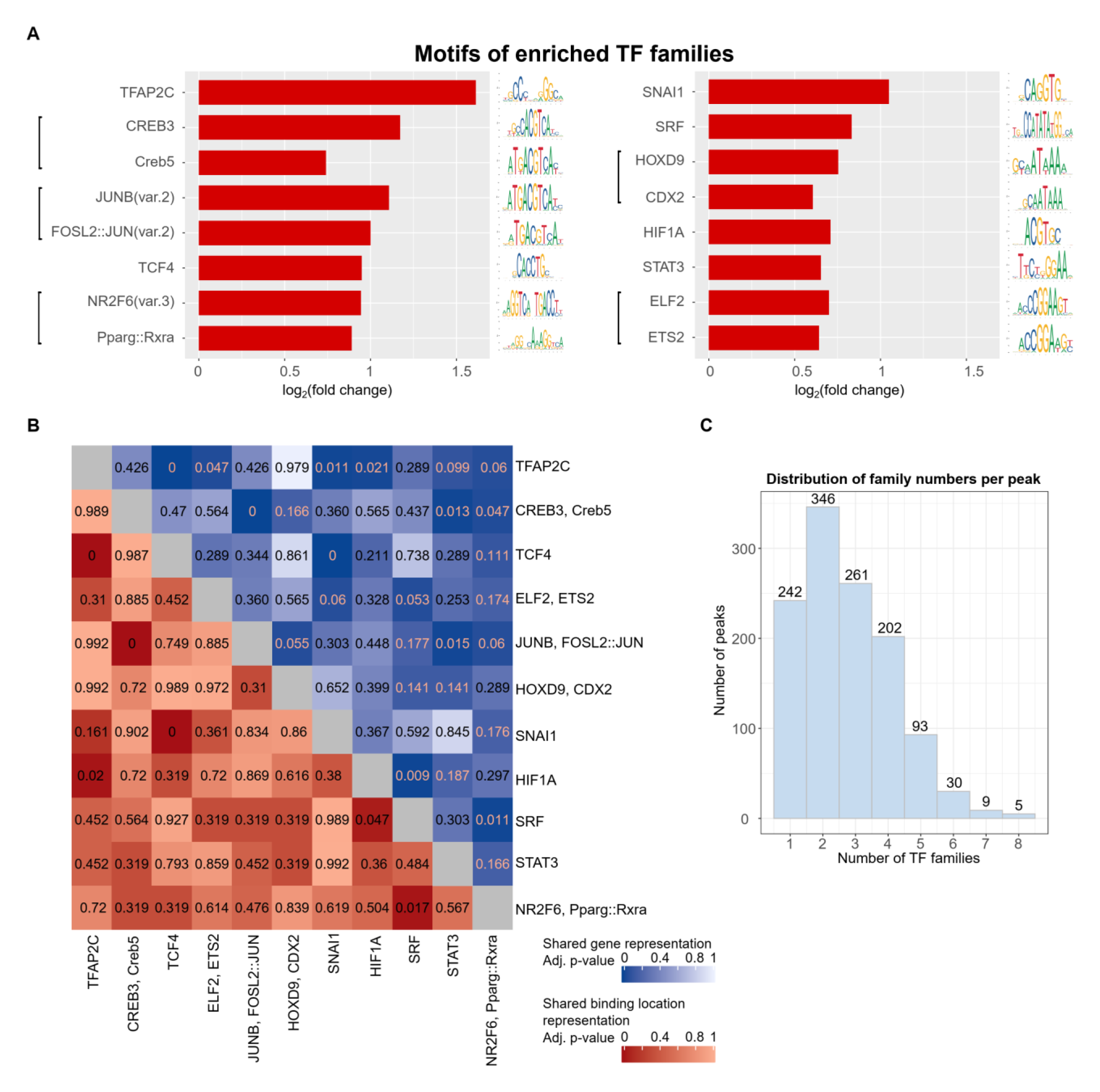
Motif analysis identifies transcription factor (TF) combinations regulating invasive trophoblast cell functions. **A**) Representative motifs for enriched TF families found in common open chromatin peaks. Motifs for the top two most highly expressed TFs in each family are shown. In case multiple motifs are enriched that correspond to the same TF, motifs with the highest fold change are shown. See the mapping of motifs to TFs in **Table S3**. A motif is considered enriched if its hypergeometric adjusted p-value is ≤0.05 and fold change ≥1.5. The p-values were adjusted with the Benjamini-Hochberg procedure. Only motifs corresponding to genes with expression level ≥0.5 at both gd 15.5 and gd 19.5 were used in the downstream analysis. **B**) Heatmap of hypergeometric adjusted (**adj.**) p-values showing that some TF family pairs share a significant number of target genes and binding locations. The p-values were adjusted with the Benjamini-Hochberg procedure. Representative motif names (as in **A**) were used for TF family names. Significance level was 0.05. Blue scale: adj. p-values when testing for significance of shared genes; dark red scale: adj. p-values when testing for significance of shared binding locations. **C**) Histogram for number of TF families per open chromatin peak showing that most open chromatin peaks had at least two TF families predicted to be bound while there were some open chromatin peaks with only one TF family predicted to be bound. The x-axis shows the number of TF families per peak, and the y-axis showed the number of peaks.

To determine if TF functions could be predicted using the binding sites, we carried out functional enrichment analysis on the genes associated with peaks where the TF families’ binding sites were found. We observed four families with at least one term enriched (**Table S3**), two of which were enriched for important invasive trophoblast functions: “NR2F6, Pparg::Rxra” (*Thyroid hormone receptor-related factors – RXR-related receptors* family*, Nuclear receptors with C4 zinc fingers* class) enriched for “positive regulation of cell migration” and “vasculature development”; and “TFAP2C” (*AP-2* family*, Basic helix-span-helix factors* class) enriched for “cell-cell adhesion”, “positive regulation of cell motility”, and “vasculature development”. Many of these observed terms agree with previous findings about roles of the families in trophoblast cell functions [44,45,52].

Next, we investigated which TF families were associated with the same target genes. We observed multiple pairs of TF families that shared a significant number of overlapping target genes, such as: “TCF4” (*E2A-related factors* family*, Basic helix-loop-helix factors* class) and “SNAI1” (*More than 3 adjacent zinc finger factors* family, *C2H2 zinc finger factors* class) (adjusted p-value=3.64e-55); “JUNB, FOSL2::JUN” (*FOS-related factors – JUN-related factors* family, *Basic leucine zipper factors* class) and “CREB3, Creb5” (*CREB-related factors* family, *Basic leucine zipper factors* class) (adjusted p-value=9.35e-41); and “TFAP2C” (*AP-2* family*, Basic helix-span-helix factors* class) and “TCF4” (*E2A-related factors* family*, Basic helix-loop-helix factors* class) (adjusted p-value=7.88e-06) (**Fig. 3B**, blue scale). Overall, this analysis highlights TF families that share common target genes.

We also checked if TF family pairs occurred in the same peaks more than expected by chance. We found six pairs of TF families significantly over-represented together, including: “TCF4” (*E2A-related factors*) and “SNAI1” (*More than 3 adjacent zinc finger factors* family) (adjusted p-value=3.09e-58), “CREB3, Creb5” (*CREB-related factors* family) and “JUNB, FOSL2::JUN” (*FOS-related factors – JUN-related factors* family) (adjusted p-value=1.36e-45), and “TFAP2C” (*AP-2* family) and “TCF4” (*E2A-related factors* family) (adjusted p-value=4.67e-04) (**Fig. 3B**, dark red scale). Each TF family that is part of the over-represented pairs has been individually connected to the regulation of trophoblast cell function. For example, TCF4 and SNAI1 are regulators of trophoblast cell differentiation and motility [53] and trophoblast invasion [54], respectively. Moreover, most of the peaks were bound by at least two TF families (**Fig. 3C**). This analysis suggested that TF families can bind in the same locations to interact and regulate cell type-specific functions. TFs can also bind individually to act in their regulatory roles.

### Identification of conserved, invasive trophoblast cell-specific regulatory regions using network analysis

To predict distal elements and TFs associated with genes defining invasive trophoblast cell clusters at both gestation days, we created a TF-gene network. To establish the network we compiled several datasets: i) rat invasive trophoblast cell common peaks that overlapped with accessible regions in EVT cells (conserved common peaks), ii) motifs enriched within these regions, and iii) conserved genes that exhibited invasive trophoblast cell-specific expression, according to the scRNA-seq analysis [20], at both gd 15.5 and 19.5 [20]. The resulting network had 11 source nodes, corresponding to 11 TF families, and 34 target genes (**Fig. 4A, Table S4**).

**Fig. 4.**
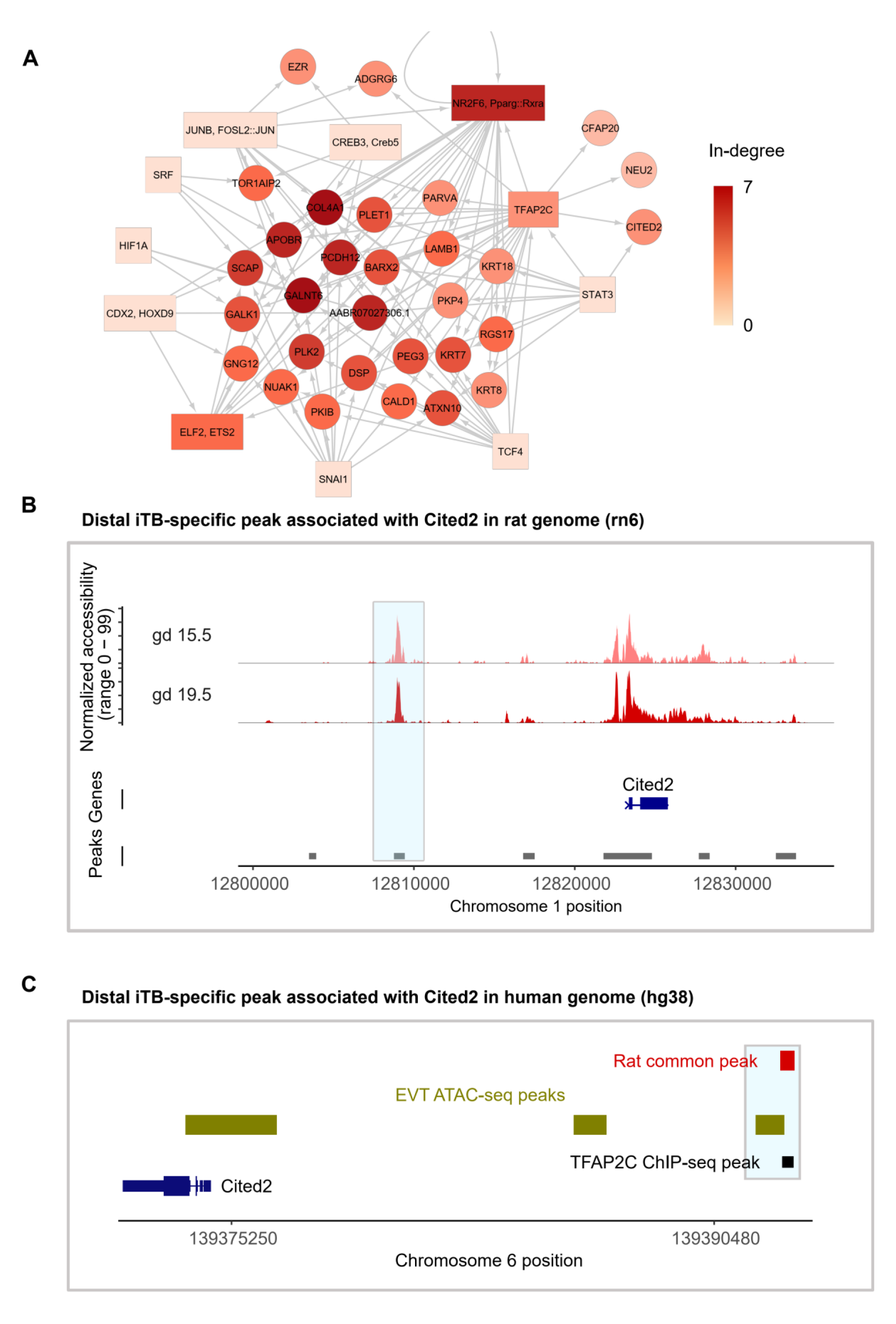
Network analysis predicts candidate genes and their distal regulatory elements that govern invasive trophoblast cell functions. **A**) Analysis of a network of TF families and target genes highlighting candidate genes and their distal regulatory elements underlying invasive trophoblast cell functions. Rectangular nodes: TF families with representative motif names (as in **Fig. 3A**). Round nodes: target genes. Color: the darker the color, the higher the node in-degree centrality. Directed edges mean peaks with the predicted TF families were associated to the target genes. **B**) Chromatin accessibility tracks of a candidate invasive trophoblast (**iTB**) cell-specific distal element associated with the *Cited2* gene in the rat genome (rn6). A region of interest was highlighted in light blue. **C**) Locations of the candidate region, ATAC-seq peaks in EVT cells and TFAP2C ChIP-seq peaks in the human genome (hg38). A region of interest was highlighted in light blue.

In this network, there are multiple genes with high in-degree centrality (≥5), meaning the genes were associated with invasive trophoblast cell-specific peaks predicted to be bound by TFs connected to ≥5 TF families. These genes were *Plk2* (linked with five TFs), *Scap* (linked with five TFs), *AABR07027306.1* (*PHACTR1* human ortholog, linked with six TFs), *Pcdh12* (linked with six TFs), *Galnt6* (linked with six TFs), and *Col4a1* (linked with seven TFs) (**Fig. 4A**). *Pcdh12*, *Plk2, Scap*, and *Col4a1* have previously been linked to the regulation of embryonic and placental development [55–61]. Although, *Phactr1* and *Galnt6* have not been directly implicated in trophoblast cell biology, they have been shown to regulate migration and invasion of cancer cells [62–64]. Further analysis of the involvement of these genes in the regulation of the invasive trophoblast cell lineage is merited. Regulatory elements and the enriched motifs associated with these genes as well as all other target genes in the network can be found in **Table S4**.

Moreover, other target genes in the network and their distal elements could also be important for regulating invasive trophoblast cell functions. For example, *Cited2*, a gene required for trophoblast cell differentiation, placental development, and regulation of invasive trophoblast/EVT cells [23,47,48,65], was predicted to be regulated by a distal peak where TFAP2C and STAT3 motifs were found (**Fig. 4A and B**). This peak (*chr1:12808761-12809434*) also overlapped with a TFAP2C ChIP-seq peak from differentiated mouse TS cells [34] (**Fig. 4C**), suggesting that it may be bound *in vivo*.

Together, the target genes, regulatory elements, and TFs we identified will be candidates for future experiments to interrogate gene regulatory networks controlling invasive trophoblast cells.

## DISCUSSION

The invasive trophoblast cell lineage is an evolutionary adaptation facilitating viviparity in mammals possessing hemochorial placentation [66]. Invasive trophoblast cells acquire migratory behavior, penetrate the uterine parenchyma, and serve a transformative role on cellular constituents ensuring a successful pregnancy outcome [3,4,7]. The root cause of many obstetric complications is predicted to be a failure in invasive trophoblast cell-guided uterine transformation [8,9]. Surprisingly, existing knowledge of gene regulatory networks controlling development and function of the invasive trophoblast cell lineage is modest. In this report, we sought to provide new insights into the regulation of the invasive trophoblast cell lineage. Our efforts focused on the rat, a species possessing deep intrauterine trophoblast cell invasion with similarities to human placentation and amenable to testing hypotheses pertaining to the invasive trophoblast cell lineage *in vivo* [12,13]. In this report, we integrated snATAC-seq and scRNA-seq [20] datasets from the rat uterine-placental interface with the goal of gaining insight into gene regulatory networks controlling the invasive trophoblast cell lineage. Chromatin accessibility profiles for each of the cellular constituents of the uterine-placental interface were determined. An in-depth analysis of invasive trophoblast cells led to the identification of invasive trophoblast cell specific genes, TFs, and TF target genes. A correlation was established between the presence of invasive trophoblast cell-specific open chromatin and gene expression. Using DNA motif binding enrichment and network analysis, we predicted TF pairs and *cis-*regulatory elements linked to invasive trophoblast cell genes. The efforts led to the recognition of conservation between rat and human invasive trophoblast cell lineages and predictions of distal regulatory elements within the invasive trophoblast cell lineage.

Our approach of relating open chromatin to gene expression profiles is not perfect. Gene regulatory regions can regulate multiple genes [67] and can be located considerable distances from the gene they regulate [68]. We observed that most open chromatin regions were distal to genes. Moreover, the open chromatin-gene association rule we used, together with the stringent requirement for conserved regulatory regions and genes, contributed to the inference of a relatively small and manageable network of TFs and target genes. This contributed to a straightforward network analysis that enabled the prediction of relevant interactions. Other computational methods such as co-accessibility analysis, which employs chromatin accessibility profiles to predict interactions of *cis*-elements [69], represents a complementary approach. Although our network construction method involved using only conserved open regions and conserved target genes, this does not negate the merits of investigating TFs and target genes inferred with species-specific elements.

Candidate TFs driving gene regulation in invasive trophoblast cells were identified through their expression in invasive trophoblast cells and through the presence of corresponding TF DNA binding motifs associated with invasive trophoblast cell specific genes. The most striking TF families linked to the invasive trophoblast cell lineage exhibit conservation in human EVT cells [39] and have been previously implicated in trophoblast cell biology [70,71]. Most interestingly, many of the invasive trophoblast cell relevant TFs are implicated in early phases of trophoblast cell lineage development or the differentiation of other trophoblast cell lineages. For example, mouse mutagenesis has demonstrated indispensable roles for *Tfap2c*, *Cdx2*, *Ets2*, and *Pparg* in trophoblast cells and placentation that precede the appearance of the invasive trophoblast cell lineage [44–46,52,72,73]. Some of these TFs were predicted to regulate the same genes based on the motif enrichment analysis, and all of these TFs had a high degree of connectivity with each other in the network we present. Previous studies have determined that TFs can work in combination to regulate trophoblast cell lineages, but different TF partnerships are implicated in the regulation of distinct processes [71,74,75]. Re-use of trophoblast lineage associated TFs in the regulation of invasive trophoblast cells is intriguing but creates experimental challenges. Future *in vivo* investigation will necessitate the establishment of conditional mutagenesis rat models specific to the invasive trophoblast cell lineage. Such efforts will be facilitated by the integration of single-nucleus chromatin accessibility and single-cell gene expression profiles reported here. Unique TF combinations at gene regulatory domains and/or the recruitment of unique sets of co-regulators may prove crucial to invasive trophoblast cell biology.

The uterine-placental tissue used in generating the snATAC-seq and scRNA-seq contains invasive trophoblast cells that have exited the placenta and entered the uterus and thus represent a differentiated cell type. We did not observe any evidence for multiple types of differentiated invasive trophoblast cell types nor did we detect evidence for invasive trophoblast cell progenitors. This latter population of progenitor cells should reside in the junctional zone of the rat placenta or the EVT cell column of the human placenta. Thus, the present analysis is biased towards characterization of a mature invasive trophoblast cell population. Consequently, the invasive trophoblast cell gene signature, including TFs, may best represent requirements for maintenance of the invasive trophoblast cell state. Comparisons of these rat invasive trophoblast cell chromatin and gene expression profiles with human EVT cell populations isolated from first trimester tissues [15–20,39] or derived from human TS cells [39,59] have some inherent limitations. Elucidation of single cell multi-omic profiles for the junctional zone will provide valuable information regarding derivation of the invasive trophoblast cell lineage and further insights into conservation of this important developmental process.

The datasets and analyses presented in this report represent a framework for constructing hypotheses relevant to establishing a gene regulatory network controlling the invasive trophoblast cell lineage. A research approach can now proceed involving identification of candidate conserved regulatory pathways, evaluating the importance of the regulators using TS cell models, and testing critical hubs within the pathways using relevant in vivo rat models.

## MATERIALS AND METHODS

### Animals

Holtzman rats were originally purchased from Envigo. Rats were maintained on a 14 h light/10 h dark cycle with open access to food and water. Timed pregnancies were obtained by mating adult males (>10 weeks of age) and adult females (8-12 weeks of age). Pregnancies were confirmed the next morning by presence of sperm in a saline vaginal lavage and defined as gd 0.5. Protocols for research with animals were approved by the University of Kansas Medical Center (**KUMC**) Animal Care and Use Committee.

### Cell isolation from tissue

Uterine-placental interface tissue (also called metrial glands) were dissected from gd 15.5 (n=3 pregnancies) and 19.5 rat placentation sites (n=3 pregnancies) as previously described [20,76] and put in ice cold Hank’s balanced salt solution (**HBSS**). Tissues were minced into fine pieces with a razor blade and digested in Dispase II (1.25 units/mL, D4693, Sigma-Aldrich), 0.4 mg/mL collagenase IV (C5138, Sigma-Aldrich), and DNase I (80 units/mL, D4513, Sigma-Aldrich) in HBSS for 30 min. Red blood cells were lysed using ACK lysis buffer (A10492-01, Thermo-Fisher), rotating at room temperature for 5 min. Samples were washed with HBSS supplemented with 2% fetal bovine serum (**FBS**, Thermo-Fisher), and DNase1 (Sigma-Aldrich) and passed through a 100 μm cell strainer (100ICS, Midwest Scientific). Following enzymatic digestion, cell debris was removed using MACS Debris Removal Solution (130-109-398, Miltenyi Biotec). Cells were then filtered through a 40 μm cell strainer (40ICS, Midwest Scientific) and cell viability was assessed, which ranged from 90 to 93%.

### Nuclei isolation, library preparation, and sequencing

Cells were isolated from gd 15.5 and 19.5 uterine-placental interface tissue as described above, and nuclei were isolated from the cell suspension according to the 10X Genomics Nuclei Isolation protocol. Briefly, cells were washed with HBSS supplemented with 2% FBS (Thermo-Fisher) and cell number determined. Approximately 500,000 cells were centrifuged, and 100 μL 10X Genomics Nuclei Isolation Lysis Buffer was added. The suspension was incubated for 3 min, then 10X Genomics Nuclei Isolation Wash Buffer was added. Cells were passed through a 40 μm cell strainer and centrifuged. Cells were resuspended in 50 μL chilled 10X Genomics Nuclei Isolation Buffer. Single nuclei were captured using the Chromium Controller into 10X barcoded gel beads. Libraries were generated using Chromium Next GEM Single Cell ATAC Library & Gel Bead Kit v1.1 (10X Genomics) and sequenced in a NovaSeq6000 sequencer at the KUMC Genome Sequencing Core.

### snATAC-seq preprocessing

Read alignment to the rat genome (Rnor 6.0, Ensembl 98 [77]), primary peak calling, and feature quantification were performed using Cell Ranger Software (version 4.0.0). Quality control steps and downstream analyses were performed using the R package Signac (version 1.1.1) [40]. Unless otherwise reported, default parameters were used. We identified accessible regions using the CallPeaks() function in Signac, which utilizes model-based analysis for ChIP-seq (**MACS**) [78]. Parameters used for the analyses were nuclei with a total number of fragments in peaks ranging from 1000 to 20000, percentage of reads in peaks >15%, and enrichment ratio at transcription start sites >1.5 (**Fig. S1**). We normalized across samples and across peaks using term frequency-inverse document frequency, which is implemented through RunTFIDF() in Seurat. We used method =3, which computes log(term frequency) × log(IDF), due to great sparsity in the feature matrix and strong count outliers (**Fig. S2**). All features are retained to perform dimension reduction with singular value decomposition (**SVD**). Normalization with term frequency-inverse document frequency followed by SVD is also known as latent semantic indexing (**LSI**) [79]. We also investigated the correlations between sequencing depth and LSI components (using the DepthCor() function) as well as ranked the LSI components using the percentage of variance (using the ElbowPlot() function). As a result, we kept LSI components 2 to 20 for gd 15.5 replicates, and LSI components 2 to 10 for gd 19.5 replicates (**Fig. S2**). Replicates for each time point were then merged using the Merge() function in Seurat.

### snATAC-seq clustering

To identify cell clusters for each time point, we utilized K-nearest neighbor (**KNN**) graphs with retained significant LSI components and the smart local moving algorithm [80], which was implemented through the Seurat functions FindNeighbors() and FindClusters(). The clusters were then visualized with uniform manifold approximation and projection (**UMAP**).

### scRNA-seq and snATAC-seq integration – label transferring

To transfer cluster labels from our corresponding scRNA-seq data, we used the FindTransferAnchors() and TransferData() functions in the Seurat package (version 4.1.0) [81]. Briefly, this process uses canonical correlation analysis for initial dimension reduction, then identifies cell neighborhoods with KNNs, and mutual nearest neighbors (**MNN**). The correspondences between cells were referred to as “anchors”. Next, the anchors were given scores and weights to eliminate incorrect correspondences and to define the association strengths between cells and anchors. Finally, anchor classification and anchor weights were used to transfer labels from scRNA-seq to snATAC-seq data.

To check the correlation between snATAC-seq and scRNA-seq profiles in each cell population, we first estimated the chromatin accessibility profiles around transcription start sites, referred to as the gene activity, using the Signac function GeneActivity(). Then Spearman correlation and its statistical significance were calculated using the R function cor.test() (*stats* package version 4.0.2 [82]).

### Analysis of cell population-specific peaks

The FindAllMarkers() function was used with cell identities transferred from scRNA-seq data and the fragment counts in peaks, to compare chromatin accessibility profiles between cell types for each gd. We used a logistic regression framework with a latent variable of the total number of fragments in peaks to account for the difference in sequencing depths. A peak is considered more accessible in a cell population (and hence specific) if it has an adjusted p-value ≤0.05 and an average log_2_(fold change) ≥log_2_(1.5).

Rat peaks were associated with the nearest gene (according to the start position) on the same chromosomes using the Signac function ClosestFeature() with the underlying genome annotation from Ensembl 98 [77]. This association rule was also used when the distance distribution of peaks to transcription start sites was calculated with the R package ChIPseeker [83]. ATAC-seq peaks in EVT cells [39] were associated to the single nearest genes with the maximum distance of 1000 kb around the TSS using GREAT (Genomic Regions Enrichment of Annotations Tool) [67]. Rat genes were mapped to their one-to-one human orthologs using gene mapping from Ensembl 98.

To assess changes in the expression level of transcripts with different numbers of associated peaks, or differences in the numbers of EVT peaks between two gene groups, we used the Wilcoxon rank sum test, implemented with the R function wilcox.test() (*stats* package version 4.0.2 [82]). To test the significance of overlap between genes with ≥3 peaks and invasive trophoblast cell markers, we used the hypergeometric test with the R function phyper() (*stats* package version 4.0.2 [82]) using options lower.tail = TRUE. In all tests, the significance level used was 0.05.

### Common peaks, peak mapping across species, and conserved common peaks

Common invasive trophoblast cell-specific peaks between the two gd were obtained using bedtools intersect (version 2.27.1) [84]. Regions between the two gd were considered common if ≥50% of the base pairs overlapped.

To compare peaks across species (rat, mouse and human), all peak sets were converted to human coordinates (hg38) using LiftOver (default settings) [85].

Bedtools intersect (version 2.27.1) [84] was used to identify conserved peaks, which were defined as peaks that overlapped with ATAC-seq peaks in EVT cells [39] by ≥1 base pair (**bp**).

### Motif analysis with common peaks

To identify enriched motifs in common peaks, we used the *Homo sapiens*, *Mus musculus* and *Rattus norvegicus* motif databases from JASPAR (version 2020) [86]. A BSgenome object for *Rattus norvegicus*, necessary to add motif information to Seurat objects, was built using the BSgenome R package (version 1.58.0) [87] and genome sequences obtained from Ensembl 98 [77]. We used the gd 19.5 coordinates of the common peak sets as input, then generated a set of 50,000 background sequences with matched length and GC content distribution using the Seurat function MatchRegionStats(). For each motif, we calculated a fold change as the percentage the motif is observed in the input sequences divided by the percentage it is observed in the background. A motif is considered enriched if its hypergeometric adjusted p-value is ≤0.05 and fold change ≥1.5. The p-values were adjusted with the Benjamini-Hochberg procedure [88].

To identify motif groups, we first mapped enriched motifs for all three organisms to their corresponding TFs using TF – motif mapping information from the JASPAR database, then retained only TFs with expression level ≥0.5 at both gd using the scRNA-seq data. Next, we grouped TFs according to their protein families, also obtained from the JASPAR database.

To compare the observed binding sites of the protein TFAP2C with previously published data from Lee et al. [34], we accessed the TFAP2C ChIP-seq data generated from differentiated TS cells through the GEO ID GSM3019344. A rat peak with TFAP2C motifs was defined to agree with mouse TFAP2C ChIP-seq peaks if they overlapped by ≥1 bp as assessed with bedtools intersect (version 2.27.1) [84]. The significance of the overlap was determined using Fisher’s exact test, with the option alternative = "greater" and a significance level of 0.05.

To carry out functional enrichment of target genes of the enriched TF families, we used Webgestalt (version 2019) [89] with the rat genome. A term was considered enriched if its FDR <0.05, enrichment rate ≥2, and number of observed genes is ≥5.

To test for over-representation of shared genes and shared binding locations, we used hypergeometric tests with the R function phyper() (*stats* package version 4.0.2 [82]) using options lower.tail = TRUE. Correction for multiple testing was carried out using the Benjamini-Hochberg procedure [88]. Significance level was set at 0.05.

### Network inferences and analyses with conserved common peaks

In our networks, an edge between a TF family and a gene means the gene is the nearest one to conserved common peaks with the enriched motifs of the family. Source nodes in the network were TF families named with representative motifs. Target genes were marker genes of the invasive trophoblast cell clusters at both gd and were conserved in EVT cells according to the scRNA-seq data [20]. The network was visualized and analyzed with Cytoscape [90].

## DATA AND RESOURCE AVAILABILITY

The snATAC-seq datasets we generated are available from the Gene Expression Omnibus website (https://www.ncbi.nlm.nih.gov/geo/GSE227943). All data generated and analyzed during this study are included in the published article and the online supporting files. All code used for the analyses are available at https://github.com/Tuteja-Lab/MetrialGland-scATAC-seq. Any additional resources generated and analyzed during the current study are available from the corresponding author upon reasonable request.

## ACKNOWLEDGEMENTS

We would like to thank the Research IT group at Iowa State University (http://researchit.las.iastate.edu) for providing servers and IT support, and members of the Tuteja and Soares laboratories for their valuable discussions.

## FUNDING

Supported by an NIH National Research Service, HD104495 (RLS), NIH grants: HD020676 (MJS), ES029280 (MJS), HD099638 (MJS), HD104033 (MJS, GT), HD105734 (MJS), HD096083 (GT) and the Sosland Foundation (MJS). Geetu Tuteja is a Pew Scholar in the Biomedical Sciences, supported by The Pew Charitable Trusts. The views expressed are those of the author(s) and do not necessarily reflect the views of the funding agencies.

## DECLARATION OF INTERESTS

The authors declare no competing interests.

---

**Fig. S1.**
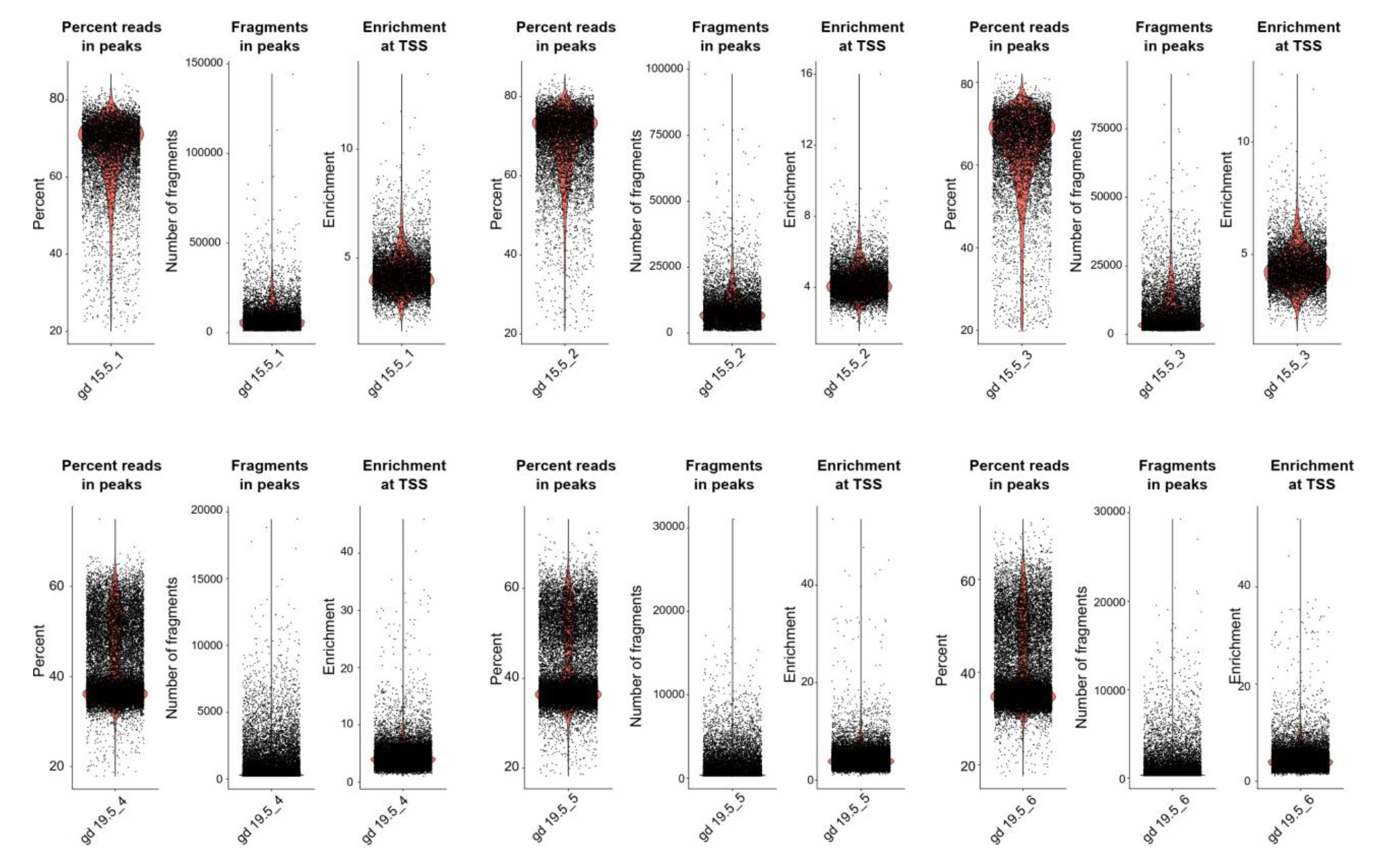
Quality control of single-nucleus ATAC sequencing (snATAC-seq) data. Violin plots showing distributions of percent of reads in peaks, numbers of fragments in peaks, and chromatin accessibility enrichment at transcription start sites (**TSS**). Nuclei with the percent of reads in peaks >15%, numbers of fragments in peaks in the range from 1000 to 20000, and chromatin accessibility enrichment >1.5 were retained.

**Fig. S2.**
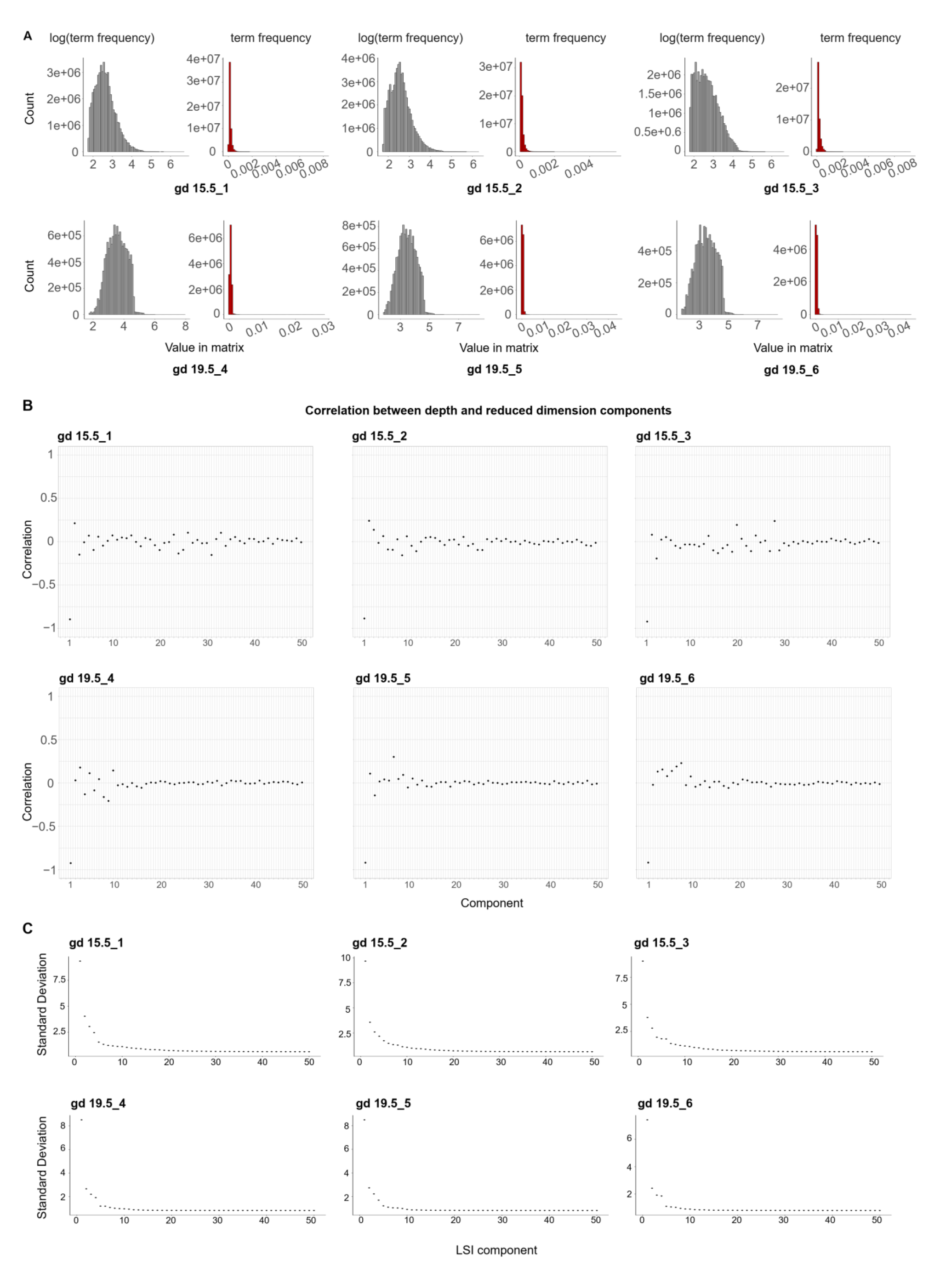
Processing of single-nucleus ATAC sequencing (snATAC-seq) data. **A**) Histogram of chromatin accessibility count matrices showing a trend of skewness. As a result, we used method =3 which computes log(term frequency) × log(IDF) for term frequency inverse document frequency normalization. Abbreviations: log(term frequency), method where log(term frequency) × log(IDF) is calculated; term frequency, method where log(term frequency × IDF) is calculated. **B**) Correlation between library depth and reduced dimension components showing that the first component across replicates were highly correlated with library depth. Therefore, the first component was excluded in the analyses. **C**) Elbow plots showing the amount of standard deviation each latent semantic indexing (**LSI**) component represented. The first 20 components of gd 15.5 samples, and 10 components of gd 19.5 samples, captured most of the variation in the data.

**Fig. S3.**
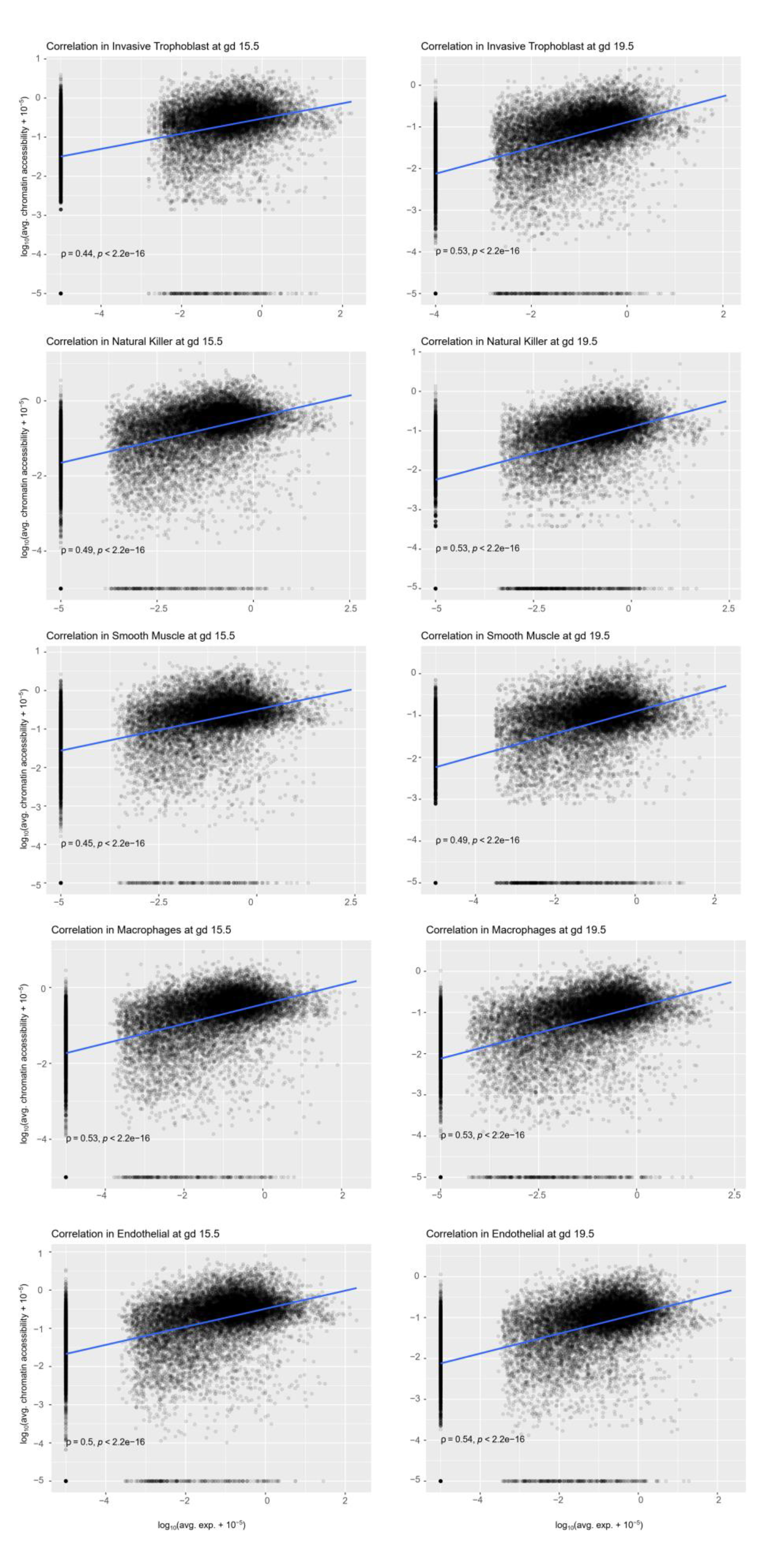
Correlation between gene expression and predicted gene activity using chromatin accessibility profiles. Scatter plots showing Spearman correlations between gene expression and predicted gene activity using chromatin accessibility profiles. At both gestation days, gene expression (x-axis) and predicted gene activity (y-axis) were moderately but significantly correlated for each of the cell populations: invasive trophoblast, natural killer, smooth muscle, macrophage and endothelial cells.

**Fig. S4.**
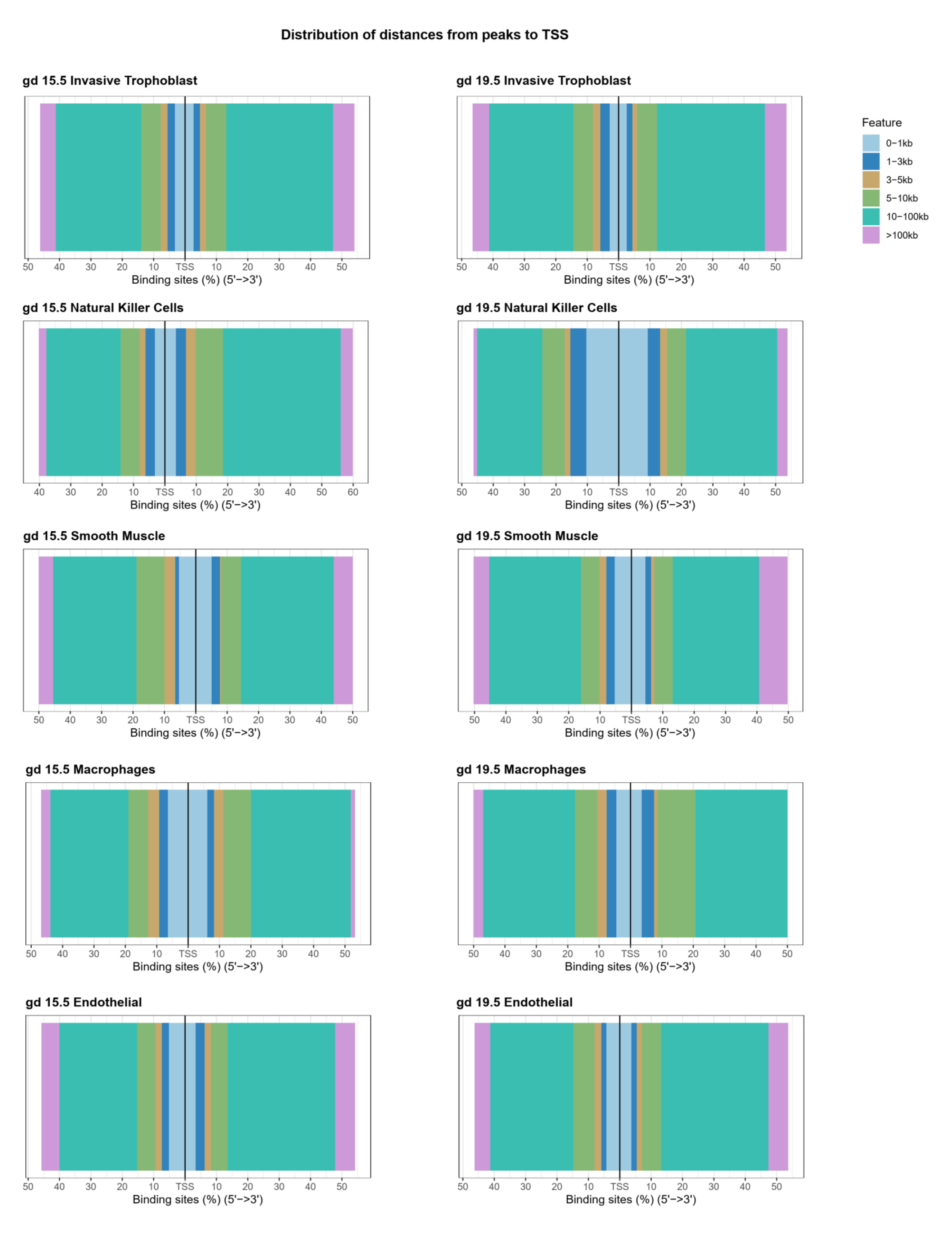
Distribution of distances between open regions and transcription start sites (TSS). Stack bar plots showing that cell type-specific open chromatin peaks were most frequently distal to the TSS.

**Fig. S5.**
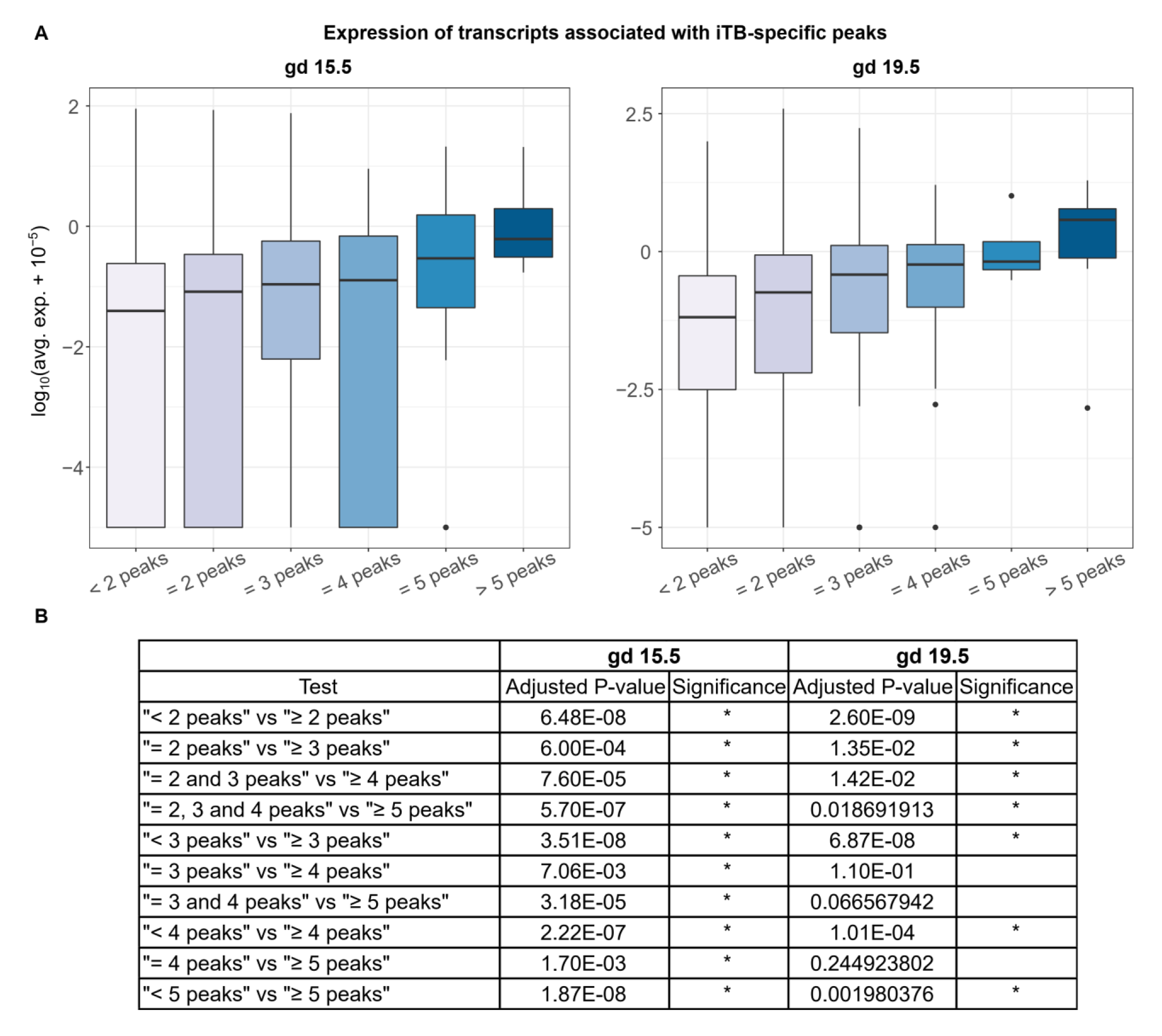
Analysis of the relationship between gene expression and the number of peaks associated with a gene. **A**) Boxplots of transcript expression associated with invasive trophoblast (**iTB**) cell-specific peaks. Expression was plotted in a log_10_(average expression + 10^-5^) scale. **B**) Adjusted p-values reported when comparing transcript expression profiles in different groups. * indicates the difference is significant. Statistical analyses were performed using Wilcoxon rank sum test at a significance level of 0.05.

## Notes

### Competing Interest Statement

The authors have declared no competing interest.

